# Targeting interactions between the Galectin-3 intrinsically disordered and structured domains based on long time-scale accelerated molecular dynamics

**DOI:** 10.1101/2021.09.27.461147

**Authors:** Supriyo Bhattacharya, Mingfeng Zhang, Weidong Hu, Tong Qi, Nora Heisterkamp

## Abstract

Intrinsically disordered regions (IDRs) are common and important functional domains in many proteins. However, IDRs are difficult to target for drug development due to the lack of defined structures which would facilitate the identification of possible drug-binding pockets. Galectin-3 is a carbohydrate-binding protein of which overexpression has been implicated in a wide variety of disorders including cancer and inflammation. Apart from its carbohydrate recognition/binding domain (CRD), Galectin-3 also contains a functionally important disordered N-terminal domain (NTD) that contacts the C-terminal domain (CTD) and could be a target for drug development. To overcome challenges involved in inhibitor design due to lack of structure and the highly dynamic nature of the NTD, we used a novel protocol combining nuclear magnetic resonance data from recombinant Galectin-3 with accelerated molecular dynamics (MD) simulations. This approach identified a pocket in the CTD with which the NTD makes frequent contact. In accordance with this model, mutation of residues L131 and L203 in this pocket caused loss of Galectin-3 agglutination ability, signifying the functional relevance of the cavity. *In*-*silico* screening was used to design candidate inhibitory peptides targeting the newly discovered cavity and experimental testing of only 3 of these yielded one peptide that inhibits the agglutination promoted by wild type Galectin-3. NMR experiments further confirmed that this peptide indeed binds to a cavity in the CTD not within the actual CRD. Our results show that it is possible to apply a combination of MD simulations and NMR experiments to precisely predict the binding interface of a disordered domain with a structured domain, and furthermore use this predicted interface for designing inhibitors. This procedure can be potentially extended to many other targets in which similar IDR interactions play a vital functional role.

## Introduction

Many multifunctional proteins contain one or more domains with no clearly-defined 3-dimensional structure (1). Although such intrinsically disordered regions (IDRs) are generally small [less than 100 amino acids] they are surprisingly abundant and have important functions in the proteins that contain them: Afafasyeva et al (2), who analyzed such structures, identified 6600 human proteins containing IDRs. The lack of a higher order structure in IDRs allows such domains to be extremely flexible, and most IDR-containing proteins are known to functionally engage in protein and RNA/DNA interactions (2).

Galectin-3 can be described as a carbohydrate-binding protein but this does not adequately capture its highly diverse cellular roles: it has been recovered at many different subcellular locations including the nucleus, the cytoplasm [at the ER-mitochondrial interface (3); in spindle poles (4) associated with lysosomes and autosomes (5)]; in membrane-less cytoplasmic ribonucleotide-protein (RNP) particles (6, 7) as well as bound to the cell surface, and secreted into the extracellular space including peripheral blood. More than 300 proteins can form complexes with Galectin-3 in hematopoietic stem cells and peripheral blood mononuclear cells (8), and Galectin-3 has been implicated in numerous pathologies ranging from heart disease and diabetes to cancer (9, 10).

The N-terminal end of Galectin-3 contains an IDR of around 80 amino acids, with the C-terminal domain (CTD) consisting of two ‘faces’. The S-face is the moiety that recognizes and binds to specific glycoproteins and includes the carbohydrate-recognition/binding domain (CRD). The function of the CRD has been studied in most detail on the surface of cells, which are covered by a dense layer of carbohydrate-containing biomolecules including glycoproteins and glycolipids. At that location, extracellular Galectin-3 regulates signal transduction strength of glycoprotein receptors through its multimerization and crosslinking activity, resulting in intermolecular and intercellular lattice complex formation (11).

Increased Galectin-3 expression correlates with many different disease states including inflammation and cancer, but a direct cause-effect relationship has also been demonstrated for some of these using knockout models. Thus, the ability to inhibit the protein is viewed as an important goal with the ultimate objective to therapeutically target Galectin-3 in different diseases (12, 13). To this end, efforts have mainly focused on the CRD: because the structure of the CTD has been determined, and the interactions of the CRD with glycans have been well-described, many carbomimetics that will interfere with the ability of Galectin-3 to bind to glycoprotein targets have been reported (14, 15). TD139 is a Galectin-3 inhibitor in this category (16) that is being tested as an inhaled drug in clinical trials for idiopathic fibrosis (17). However, some of such compounds may have unfavorable pharmacokinetic properties (18) and, as reviewed in (19), there are currently few examples of glycan-directed therapies that have transitioned to clinical use. This may be also due to challenges relating to shallow solvent-exposed binding surfaces, lack of many hydrophobic residues for ligand contact and low residence time of the bound inhibitors when lectins bind to their carbohydrates.

The N-terminal domain of Galectin-3 also appears to have a critically important contribution to its function (20) and was recently shown to mediate protein multimerization (21). Removal of this domain yields a CTD Galectin-3 protein with dominant negative activity (22–24). The CTD of Galectin-3 also contains a domain that is not the main site of direct carbohydrate recognition/binding called the F-face. Ippel et al (25) showed that the NTD interacts transiently with the CTD F-face and characterized this interaction in more detail. Moreover, Lin et al (26) reported that the disordered N-terminal domain including amino acids 20-100 forms a fuzzy complex with β-strand regions of the F-face. Importantly, the NTD mediates liquid-liquid phase separation of Galectin-3 (21, 27) which could explain its contribution to forming membrane-less structures such as cytoplasmic RNP.

The NMR-based chemical shift differences measured by Ippel et al (25) between full-length and CTD-only Galectin-3 provided information about the dynamics, but these are averaged values and do not inform on individual structures. However, it is likely that the Galectin-3 IDR will adopt an ensemble of structurally diverse conformations, that transition in the picosecond to millisecond timescale under physiological conditions and which poses serious challenges to the application of computational methods. Here, we have approached the general problem of IDR characterization using accelerated molecular dynamics (AMD) combined with existing structural data of Galectin-3 to predict the binding interface of the CTD with the IDR. The CTD binding/N-terminal interface, as observed in the AMD simulations, includes a diverse ensemble of structures in which multiple amino acid motifs between residues 20-100 of Galectin-3 engage with the CTD. We show that these structures collectively explain the NMR data from Ippel et al (25) and agree with the fuzzy complex model of IDR interaction. *In*-*silico* designed peptides based on the interacting N-terminal motifs were then used to validate the model predicted by AMD. The process described here could be used to economically target other IDR interactions with proteins or protein domains with defined structures.

## Materials and Methods

### Molecular modeling

We retrieved the human Galectin-3 CTD crystal structure from the PDB Databank (PDB ID: 6FOF) (28). The NTD was added as a random chain using Modeller (29). The full- length structure was subjected to 100 ns of MD simulation in an implicit solvent environment (30). The protein conformations were then clustered by backbone RMSD and the mean radius of gyration was calculated for each cluster. We selected a representative structure from the cluster of which the mean radius of gyration was closest to the experimentally measured one for Galectin-3 (26). This structure was used as the starting conformation for the AMD simulations. The starting structure for the AMD was solvated in explicit water and ions were added to neutralize the net charge. The system was parameterized using the a99sb-disp force field, which has been shown to perform well with both folded and disordered proteins (31). Since Galectin-3 consists of both a folded and a disordered domain, this force field is a suitable choice. Further, hydrogen mass repartitioning was implemented in order to use a 4 fs timestep (32). The system was first heated at constant volume from 0K to 310K over 30 ns with harmonic restraints applied to the protein heavy atoms. Then the system was equilibrated for 50 ns in the NPT ensemble, while the heavy atom restraints were gradually reduced to zero. Finally, the system was equilibrated for a further 50 ns unrestrained.

Five independent AMD simulations at 310K (NPT ensemble), each lasting for 250 ns were performed using the GPU accelerated AMBER software package (33). The Galectin-3 NTD conformations resulting from the five simulations were clustered by backbone dihedral RMSD and for each cluster, the average number of NTD-CTD contacts was determined. Two residues of which the Cα atoms were within 8.5 Å were defined as a contact. We also calculated the average per residue chemical shift difference (Δ*δ*) for each cluster using the software SHIFTX2 (34). For calculating Δ*δ*, we calculated the chemical shifts for the full-length Galectin-3 and those of the CTD domain only by truncating the NTD region. The Δ*δ* was then obtained as the difference between the two shifts as in Ippel et al (25). Finally, the RMSD between the calculated and experimental Δ*δ* was determined for each cluster and plotted against the number of NTD-CTD contacts (**Fig. 1B**). The clusters with the lowest Δ*δ* RMSD as well as with average number of NTD-CTD contacts greater than five (circled clusters in **Fig. 1B**) were combined to obtain a conformational ensemble with significant NTD-CTD interactions, and that is in agreement with experimental NMR data.

**Fig. 1.**
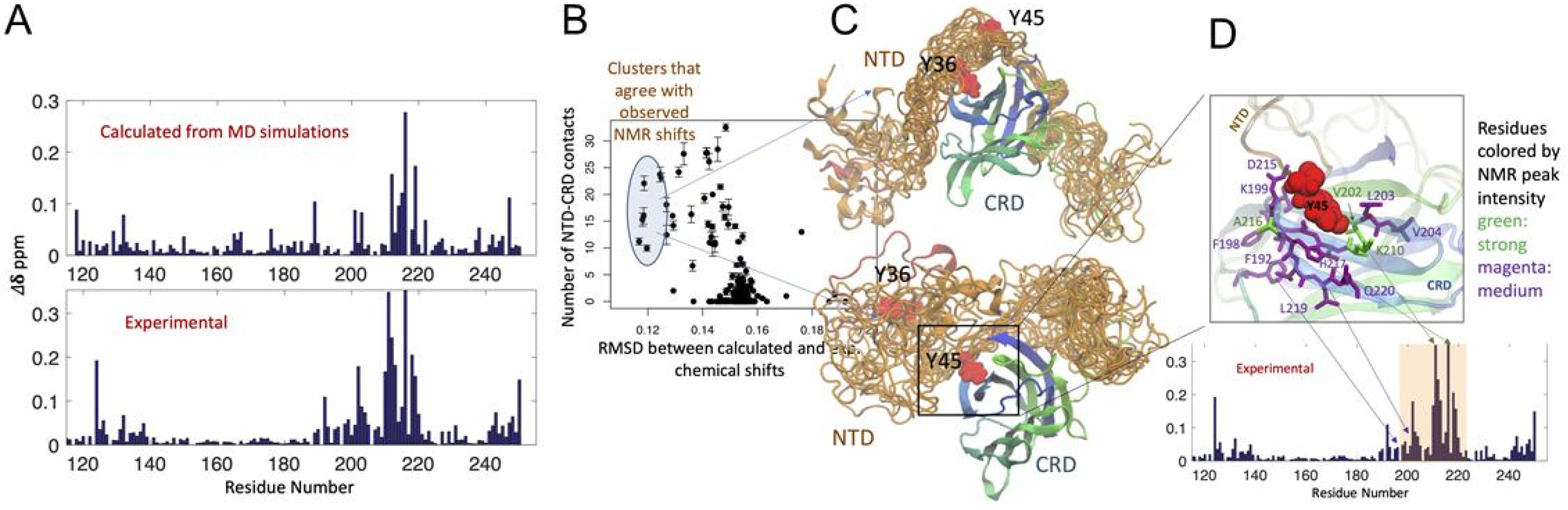
NMR-guided galectin-3 conformer generation. **(A)** comparison of the experimental chemical shift differences to those calculated from the AMD structural ensemble after filtering using the chemical shift data; **(B)** conformations from the AMD simulations were clustered by structural similarity; for each cluster, the number of NTD-CRD contacts and root mean square deviation from experimental chemical shift differences are plotted against each other; clusters that show high NTD-CRD contacts and low RMSD with experimental chemical shift differences are circled; **(C)** the two major NTD structural ensembles that agree with the experimental NMR data are shown; NTD residues that form major contacts with CRD in these ensembles (Y36 and Y45) are highlighted; **(D)** expanded view of the interface between the NTD and the CRD showing major CRD residues in contact with the NTD; the residues are colored by their peak intensities in the experimental chemical shift difference plot shown below; green: strong intensity, magenta: medium intensity.

### Bayesian maximum entropy method

The details of the BME approach is described in (35). Briefly, the weights for the AMD derived protein ensemble ([*w*_l_ … *w*_*n*_], *n*: total number of conformations) were obtained by minimizing the cost function 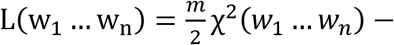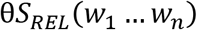, where 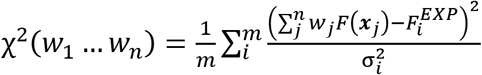 is the agreement between observed and experimental CSDs and and 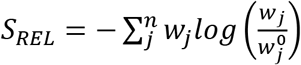 is the entropy relative to starting weights. Here, *x*_*j*_ denotes the set of protein coordinates for the *j*^*th*^ conformation, *F*(*x*_*j*_) represents the calculated CSDs using SHIFTX2 and 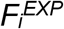 is the CSD for the i^th^ residue. *m* is the number of residues for which experimental CSDs are available. Initially, all conformations were assigned the same weight *W*_*0*_, where *W*_*0*_ = 1/*n*· σ_*i*_ denotes the uncertainty of SHIFTX2 in calculating the ^1^H and ^15^N CSDs from structure and are obtained from (34). θ is an adjustable parameter that determines the tradeoff between the entropy and the agreement with experiments. The optimal value of θ was determined by performing the optimization for different values of θ and locating the elbow of the χ^2^ vs. *log*_l0_(θ) curve, as suggested by (35). The optimization was carried out using the ‘stats’ package in R.

### Cells, culture and agglutination assay

LAX56 human pre-B ALL cells were routinely co-cultured with mitomycin-C inactivated OP9 stromal cells. These previously described primary leukemia cell grew directly out from a relapse bone marrow sample (36, 37). For agglutination assays, cells were harvested, washed once in α-MEM medium, resuspended in 10 ml X-VIVO 15 medium (Lonza) and incubated at 37°C for 24h remove the Galectin-3 produced by OP9 stromal cells. For the assay, ALL cells were resuspended in X-VIVO 15 medium at a concentration of 1 × 10^6^/ml and seeded at 2×10^5^/200 μl cells into wells. GST or GST-Galectin-3 (150 μg/ml, Fig. 4; 25 μg/ml Fig. 5) were added in 300 μl X-VIVO 15 medium to wells. Peptides, if included, were preincubated for 5 minutes with the recombinant proteins and added at different concentrations as indicated in the figures. TD139 was purchased from MedChemExpress and used at 100 μM. Phase contrast images were taken after 1-2 hours. Agglutination was defined as aggregates containing >10 cells per cluster. 5-13 images from different areas were taken and evaluated for cell clusters per condition. Biological data were graphed with GraphPad Prism software (version 8.3.1). Values represent mean ±SEM of the number of aggregates scored per independent image.

### GST-Galectin-3 and mutants

Full-length human GST-Galectin-3 (hereafter named GST-Gal3) in pGEX2T was previously described (38). To generate mutants, we used Takara on-line primer design tools and a Takara In-Fusion® Snap Assembly Kit (Cat# 638945) to generate mutations according to the manufacturer’s instructions. DNAs run on agarose gels were purified using a Thermo Scientific GeneJET Gel Extraction Kit (Cat# K0691). In-Fusion reactions (Takara) were assembled and Stellar competent cells (Takara) used for transformation. All constructs were verified by DNA sequencing (Eton Bioscience, San Diego, CA).

### Galectin-3 CTD construct for NMR

The Galectin-3 C-terminal domain construct was generated using the same methods described above for the mutants. The protein includes Galectin-3 amino acids P117-I250 as well as residual attached glycine and serine residues after thrombin cleavage. Single colonies were grown overnight in LB medium, collected by centrifugation then inoculated in M9 medium with ammonium-^15^N chloride (Sigma, Cat# 299251) and grown for 3-4 hours. After induction of protein production with IPTG for an 3-4 additional hrs, cells were harvested and suspended in 1% NP40, PI, PMSF, 1 mM DTT, 50 mM Tris-HCl, pH 7.5. Cells were disrupted by sonication. GST-Galectin-3 was bound to glutathione-agarose (Genscript, Cat# L00207) overnight at 4°C. Beads were washed 4× in lysis buffer, then suspended in 50 mM Tris-HCl pH 7.5, 0.1 mM DTT and treated with 60 U thrombin/ml (Cytiva Thrombin Protease, Fisher Scientific, Cat# 45-001-320) for 16 hrs at RT. The supernatant containing Galectin-3 protein was treated with benzamidine sepharose (HiTrap Benzamidine FF, Sigma, Cat# GE17-5143-02) to bind and remove thrombin. Protein was concentrated using an Amicon 3K filter and used in 20 mM potassium phosphate buffer pH 6.8, 0.1 mM DTT for NMR. Protein concentrations were determined by BCA.

### NMR experiments and data analysis

^1^H-^15^N HSQC experiments were carried out at 30°C on a 700 MHz Bruker AvanceIII with a TXI-triple resonance cryoprobe. The 20 μM Galectin-3 CRD was prepared in 20 mM potassium phosphate buffer, pH 6.8, 0.1 mM DTT and complex with peptide-3 in different molar ratios are indicated in Fig. 6. The data were processed and analyzed using NMRPipe (39) and NMRFAM-SPARKY (40). The chemical shift perturbation (CSP) Δδ in unit of Hz was calculated using the following equation: 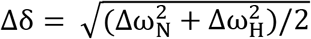. The Δω_N_ and Δω_H_ are the nitrogen and proton chemical shift difference between free ^15^N-CRD and that in the mixture with P3 peptide. Assignments are based on the Galectin-3 CRD NMR data of Ippel (25) and Umemoto (41). N-terminal domain sequences in our construct are slightly different from theirs, causing limited miss-assignment in the N-terminal domain, and ambiguity between residues 240-248 due to their close contact with the short β-strand in the slightly different N-terminal domains. Because L135 and W181 patterns differ between Ippel and Umemoto, their assignments could not be unambiguously determined. In addition, the position of T137 differs between Ippel and Umemoto, and position changes of T248 make its assignments unclear. However, none of these residues appear to be involved in the interaction with P3 peptide since their chemical shift perturbation was quite small, except for residue T248, with a CSP about 10.9 Hz and one unit of RMSD.

### Peptides

Peptides were purified by HPLC. These included peptide-1 ACE-ARAMGYPGASY-NH2, peptide-2 N-terminal acetyl -ARAFGYPIYSY-C-terminal amide and peptide-3 ACE-YYPGAYPRRYR-NH2. Peptide-4 was the Galectin-3 inhibitory peptide ANTPCGPYTHDCPVKR G3-C12 described in Zou et al (42) to target the CRD and peptide-5 the scrambled negative control peptide PTHVTCKYCPAGNRDP G3-H12s described in the same study. Neither peptide had an effect on Galectin-3-mediated agglutination (not shown).

## Results

### Derivation of the Galectin-3 NTD ensemble

An enhanced MD method called accelerated MD (AMD) uses an innovative energy rescaling method to access timescales in the order of milliseconds, that are beyond the reach of conventional MD (43). Therefore, we applied AMD to the problem of IDR conformational sampling of the Galectin-3 NTD. Starting from the initial Galectin-3 structure, where the CRD was modeled based on an existing crystal structure and the NTD was modeled as a random polymer chain, AMD was used to generate the initial conformational ensemble consisting of 50,000 NTD conformations. For each of these conformations, the corresponding chemical shifts were predicted using SHIFTX2 software, for both the full length protein as well as for the CRD alone (34). The chemical shift differences (CSDs) were then calculated according to the formula Δ*δ ppm* = [(Δ^l^*H*)^2^ + (0.25Δ^l5^*N*)^2^]^l/2^, where Δ^l5^*N* and Δ^1^*H* are the chemical shift differences of the ^15^N labeled backbone nitrogen and hydrogen atoms between full length and CTD-only Galectin-3 (**Fig. 1A, top panel**). The NTD conformations were clustered by their structural similarity (details in the methods section) and for each cluster, the root mean square deviation (RMSD) from the experimental NMR CSDs (25) were calculated. The clusters showing low CSD RMSD and a high number of NTD-CTD contacts, including a total of 1300 conformations, were selected for further processing (**Fig. 1B**). As shown in **Fig. 1A** there was an excellent agreement between the AMD-calculated and experimental CSDs within the filtered ensemble. The selected clusters are highlighted in **Fig. 1B**. By analyzing the NTD conformations that showed agreement with the experimental NMR data, two major classes of NTD-CTD contacts were identified, where of all residues, Y36 and Y45 of the NTD made the most long-term contact with the CRD, with a shallow cavity in the CTD as shown in **Fig. 1C, D**. The model predicted that the cavity would encompass candidate contacts including residues F192, F198, K199, Q201, V202, L203, V204, K210, D215, A216, H217, L219 and Q220. These residues that show close contact with the NTD in the MD ensemble also correspond to the strongest peaks in the experimental CSD profile, as shown **in Fig. 1D.**

### Bayesian maximum entropy (BME) approach uncovers diverse CTD- bound NTD conformations

We also further investigated the NTD-CRD interaction obtained from AMD using the Bayesian maximum entropy (BME) method. The details of the BME approach are given in the methods section. In brief, the BME approach tries to achieve agreement between an MD-derived ensemble and available experimental data, while maximizing the information entropy within the obtained ensemble. This leads to a conformational ensemble that maintains its diversity, while still agreeing with the experimental data. The BME approach assigns a weight to each conformation, which is proportional to its contribution to the measured experimental property.

By applying the BME approach, and using the per residue CSDs as experimental data, we calculated the weight of each NTD conformation from AMD. The highest weighted conformations were then clustered by dihedral RMSD and within each cluster, the frequencies of pairwise residue contacts between the NTD and the CRD were obtained. **Fig. 2** shows the normalized frequency of each inter-residue contact within the different clusters. Applying the BME approach, we therefore obtained a diverse ensemble, in which, apart from Y36 and Y45, multiple NTD residues have significant interactions with the CRD. The contacts where the CRD residue shows a significant peak in the experimental CSD profile are highlighted in red in the heatmap. Interestingly, we find that multiple NTD residues, notably several aromatic residues such as W22, Y101, Y41, Y45, Y54, Y70, Y79 make contact with the CRD in a way that satisfies the NMR data. Also, looking at the pairwise interactions, it appears that in many cases, multiple NTD residues interact with a single CRD residue in different conformations. Examples of such contacts include Y41 / Y45 / G47 / Q48 (NTD) → D215 (CRD), Y79 / A73 / T104 / T98 / P71 / P106 / Y89 / Y54 (NTD) → Y247 (CRD) and A100 / G112 (NTD) → T243 (CRD).

**Fig. 2.**
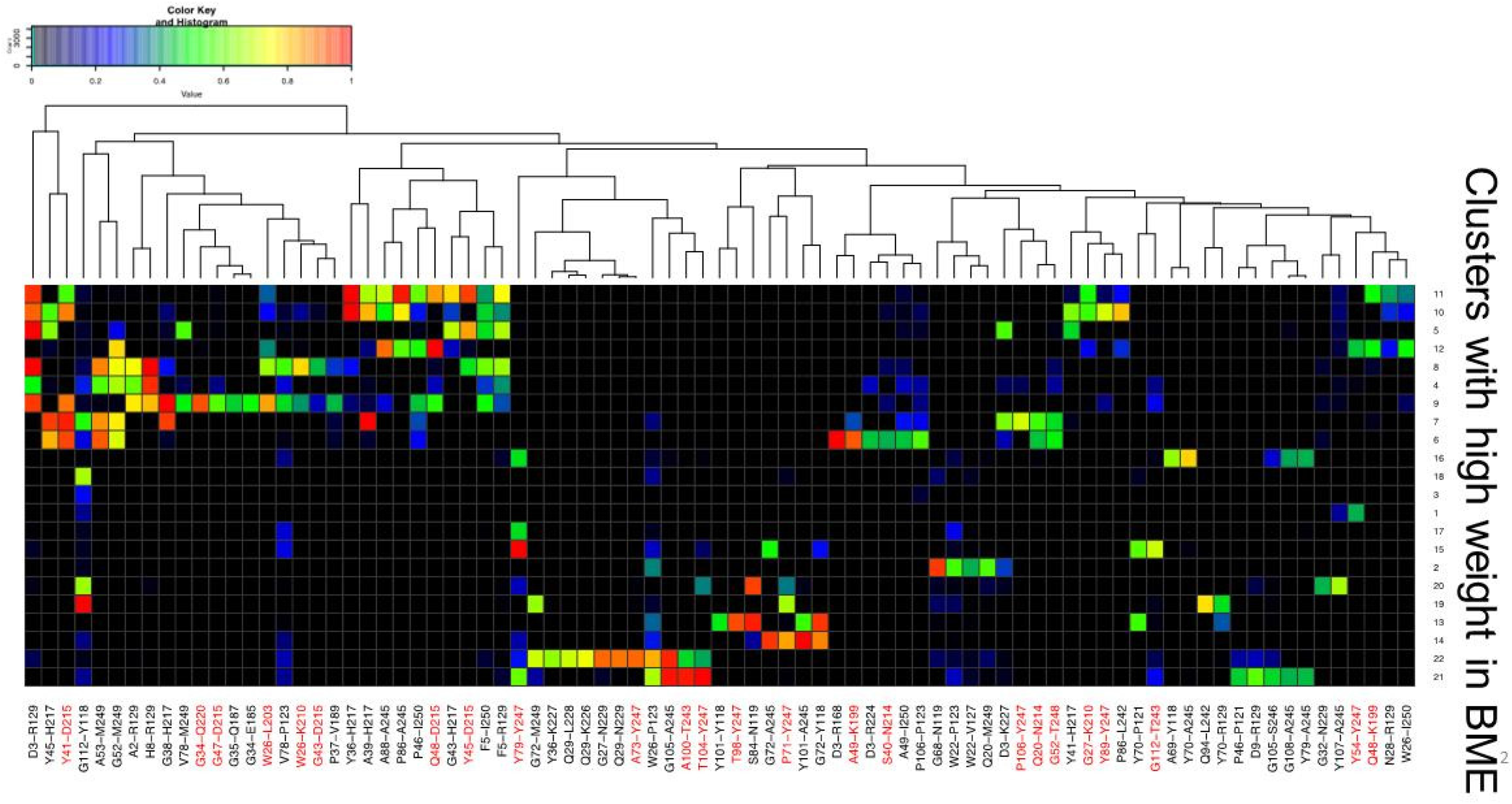
Contact heatmap of conformations with high BME weights. CTD residues with significant experimental NMR shifts are as indicated with red font below the heatmap. NTD residues making frequent CTD contacts include A2, A49, A53, A69, D3, F5, G108, G112, G43, G47, G52, G68, G72, H8, P106, P71, Q20, Q48, S84, T98, V78, W22, Y101, Y41, Y45, Y54, Y70 and Y79.

Although the BME results predicted multiple NTD-CRD contacts, the AMD-generated model pointed to specific IDR-CTD contacts that could involve a targetable pocket. We used two approaches to test this experimentally. Mutation of critical residues in that pocket could abolish binding to the IDR and the agglutination activity of Galectin-3. Also, a peptide could potentially fit in the shallow pocket and inhibit the IDR interaction. A classical test for carbohydrate-binding activity of a lectin including Galectin-3 is an agglutination assay (36, 38, 44). In this assay, recombinant Galectin-3 is tested for its ability to promote lattice formation by binding in a multivalent manner to glycoproteins located on the cell surface: when cell surface glycoprotein targets are located on different cells, carbohydrate binding combined with multimer formation causes cellular agglutination. Such an assay has widespread use for testing Galectin-3 inhibitors [e.g., (21, 42)]. Thus, we used an agglutination assay in which recombinant Galectin-3 is added to patient-derived precursor B-acute lymphoblastic leukemia cells (pre-B ALL) as a readout for Galectin-3 lattice-forming activity.

### Computational design of inhibitory peptide sequences

To design inhibitory peptides, a limited number of backbone templates were initially selected based on the ensemble of NTD conformations that showed agreement with NMR. The NTD conformations were clustered by similarity and the representative NTD conformations from the most populated clusters were selected for template design. The initial templates were obtained by retaining 5 amino acids on both sides of Y36 or Y45 in the CRD bound NTD conformations. The main steps involved in obtaining the peptide templates from the Galectin-3 NTD ensemble are shown in **Fig. 3A and 3B**. Starting from a given peptide template, each residue was systematically mutated to all 20 amino acids and an affinity score was calculated using Maestro™ software (Schrodinger LLC.), which represents the improvement in affinity of the mutant peptide over the starting NTD sequence (45). The top scoring mutations were analyzed to identify 2-3 positions in each template that were most amenable to mutagenesis. These positions were then mutated combinatorically to generate multiple double and triple mutants, and the top mutants by affinity score were analyzed for features such as strong interaction with the CTD hydrophobic cavity, low desolvation energy and sequence diversity. This step generated 8 peptide candidates, which were then subjected to 500 ns of all-atom MD simulations in an explicit water environment, to test their stability of binding to the CRD. Also, the binding free energies were calculated using the MM-PBSA method in the AMBER software package (46) (**Table 1** and **Fig. 3C**). During MD, 4/8 peptides left the CTD cavity within 300 ns and were deemed unstable (**Table 1** and **Fig. 3D**). Among the rest which remained bound and also showed strong interaction with the CRD as measured by the protein-peptide energy and number of hydrogen bonds, one peptide, Y45_cls70_Y1_M8_R9_F10_R11, was very similar in sequence to another peptide in the list and hence eliminated. The other three peptides were subjected to experimental testing (**Table 1**).

**Fig. 3.**
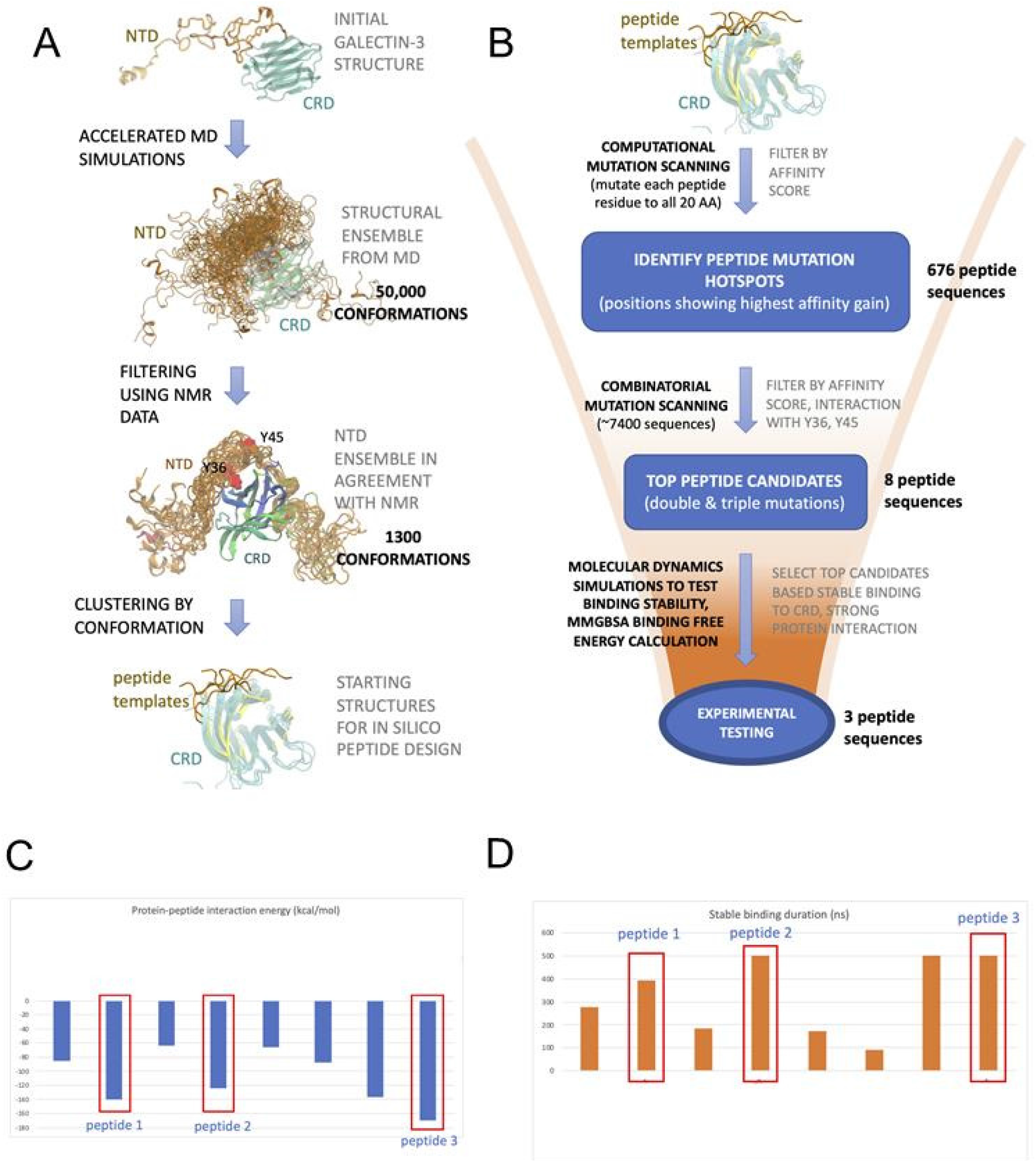
*In silico* peptide design. **(A, B)** Steps in generating the **(A)** initial peptide templates and **(B)** top peptide candidates starting from the initial templates; (C) the protein-peptide interaction energies of the top 8 candidate peptides; **(D)** duration of binding to the CRD.

**Fig. 4.**
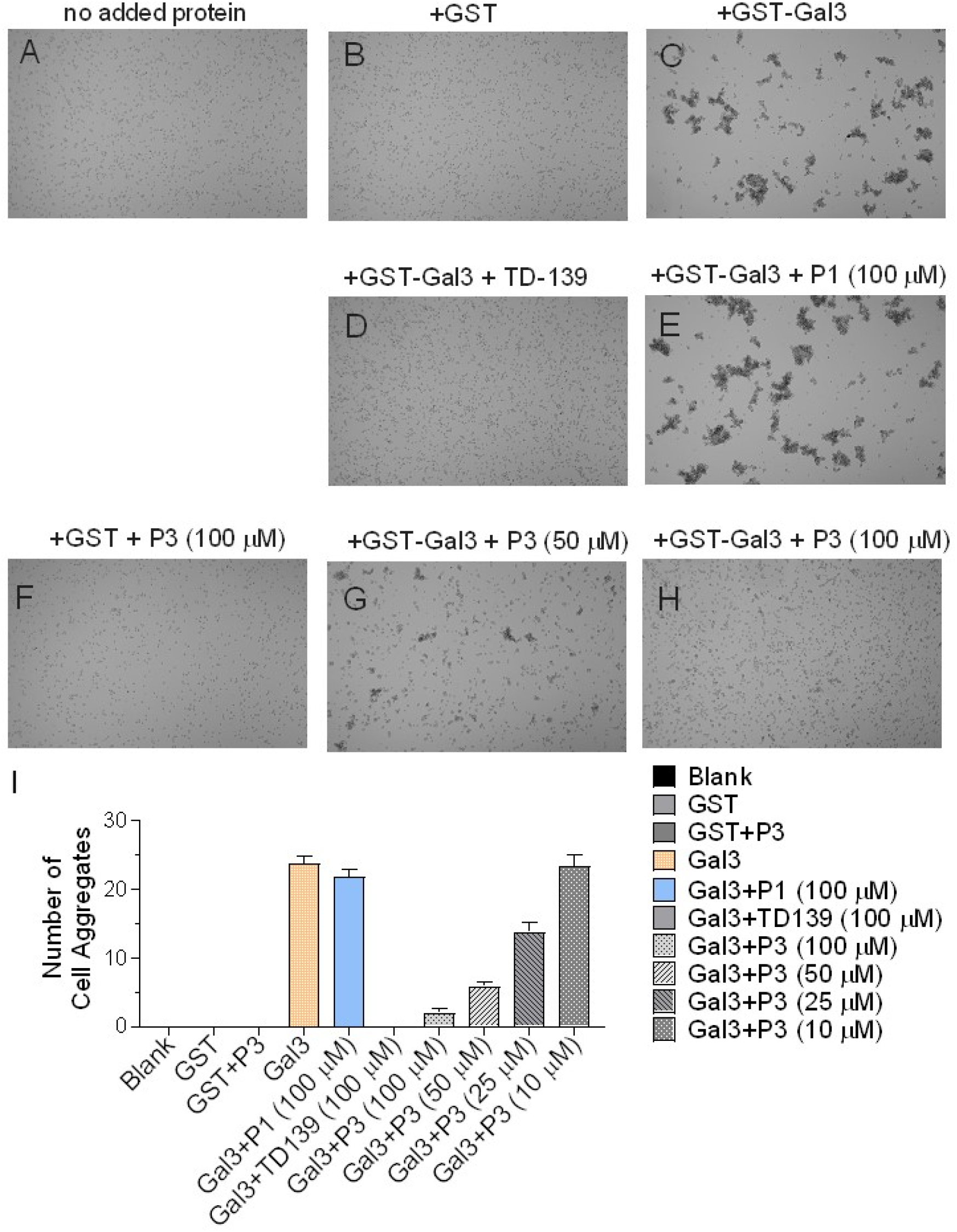
Peptide-3 inhibits agglutination of human leukemia cells mediated by Galectin-3. Representative bright field images of suspensions of pediatric pre-B acute lymphoblastic leukemia LAX56 cells. Cells were incubated for 2 hours **(A)** with no added protein **(B)** with control GST, or **(C**-**G)** with GST-Gal3. **(D)** Addition of 100 μM TD139 Galectin-3 inhibitor. **(E-H)** Peptides as indicated were added together with GST or GST-Gal3. Similar results obtained were obtained with two independently generated batches of GST-Gal3; representative images are shown. **(I)** Quantification of agglutination expressed as the number of cell aggregates. Agglutination is defined as aggregates containing >10 cells per cluster. Error bars, mean ±SEM of cell cluster counts from 5-13 images from different areas per condition.

**Fig. 5.**
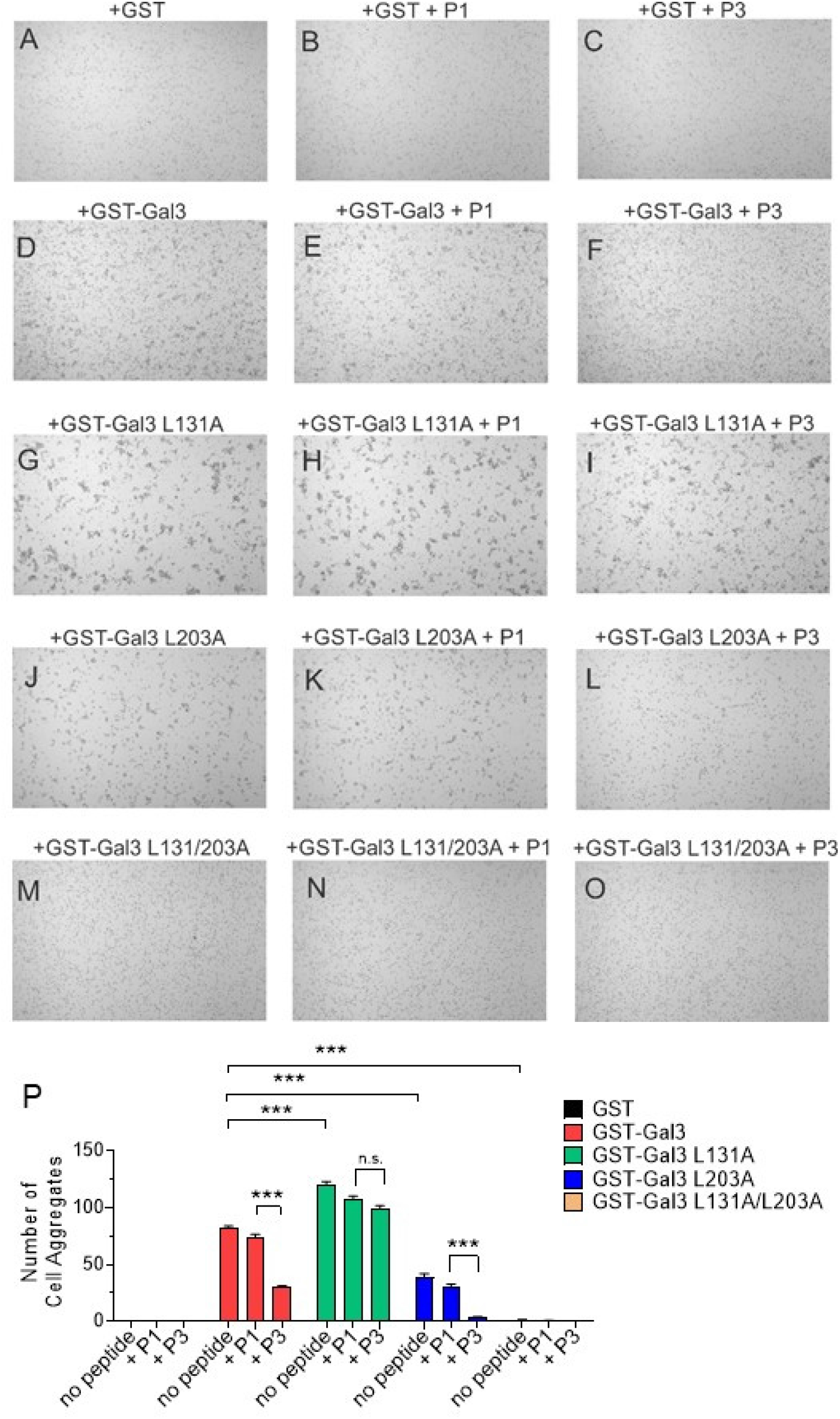
Site-directed mutagenesis of Galectin-3 shows combined L131 and L203 are essential for agglutination. **(A-O)** Bright field images of LAX56 cells incubated with the recombinant fusion proteins indicated above the panels at a concentration of 25 μg/ml. Peptides P1 or P3 added at 100 μg/ml together with the recombinant proteins are also noted above the images. **(P)** Quantitation of cellular aggregation under the indicated conditions. Error bars, mean ±SEM. P1, peptide-1; P3, peptide-3. ***p<0.001. n.s., not significant.

**Fig. 6.**
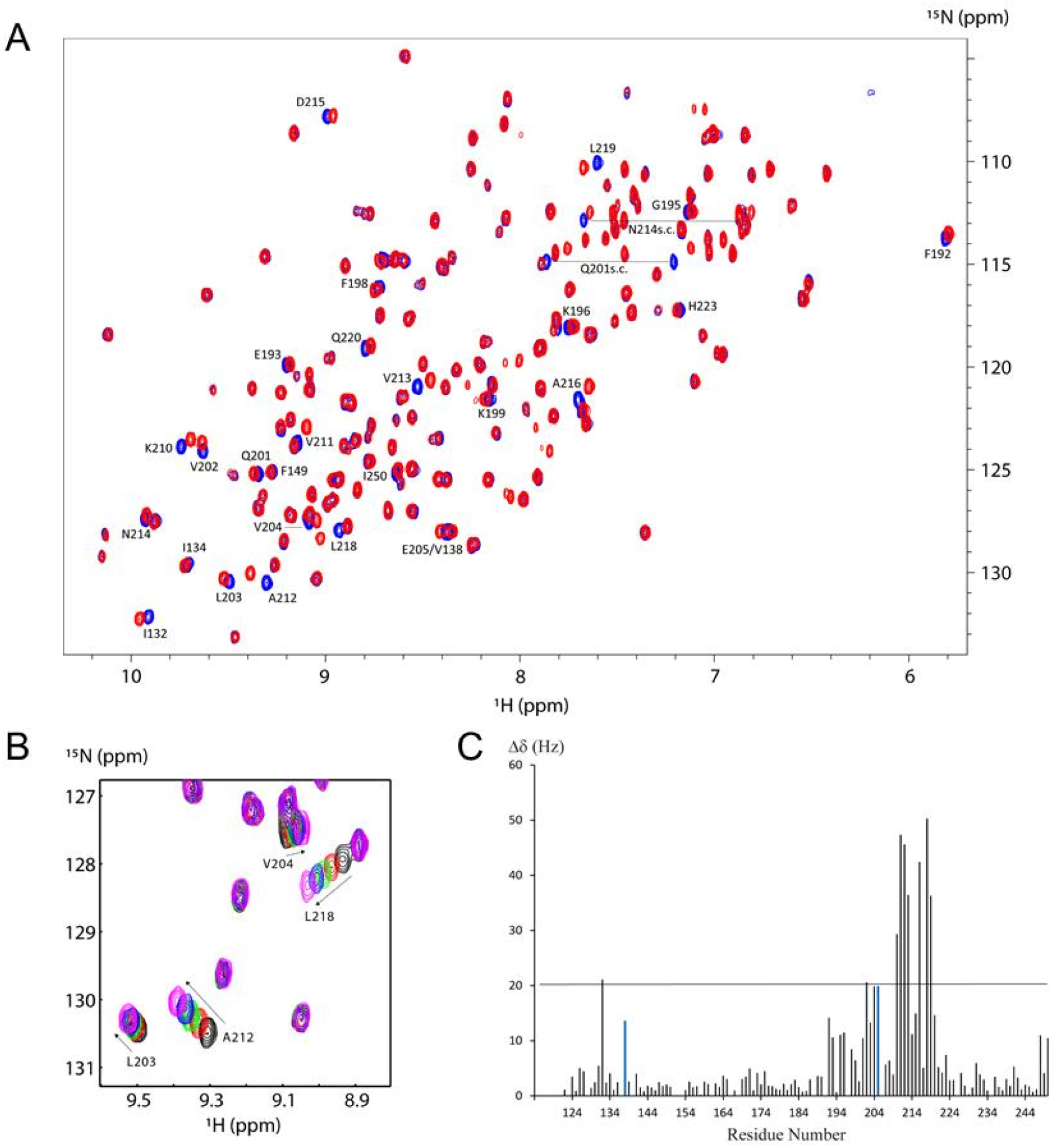
Residues in the Galectin-3 CRD domain with significant chemical shift perturbations through interaction with P3 peptide-3. **(A)** The ^1^H-^15^N HSQC spectrum in blue was acquired on free ^15^N labeled Galectin-3 CRD. The spectrum in red was acquired on the complex between ^15^N labeled CRD and peptide-3 with molar ratio of 100:1 between P3 peptide and ^15^N labeled CRD. Some residues with significant chemical shift perturbations are labeled, including two side chains from Q201 and N214. **(B)** Selected overlay of ^1^H-^15^N HSQC spectrum region of ^15^N labeled CRD versus titration of P3 peptide. Residues with notable chemical shift changes are labeled together with the cross peak moving direction, as indicated by the arrow, with increased concentration of P3 peptide. Spectrum in black is free CRD; spectra in red, green, blue and magenta are from complexes with molar ratio of 20:1, 40:1, 60:1 and 100:1 between P3 peptide and Galectin-3 CRD, respectively. **(C)** Chemical shift perturbation of ^15^N-CRD in complex with P3 peptide *versus* primary sequence of residues 117-250. The chemical shift changes between free ^15^N-CRD and in complex with 100-fold molar excess of P3 peptide are presented. The thin horizontal line indicates the limit above which values of ^15^N-CRD in complex with P3 peptide are two times the RMSD of the. Residues of V138 and E205 are color-coded in blue since they shifted apart in complex from the overlaid cross peak in free ^15^N-CRD, and their CSP values could be swapped.

**Table 1.** Binding properties for eight candidate peptides calculated from all-atom MD simulations. The peptides in bold are the best binders according to their duration of binding to the CRD, number of protein-peptide hydrogen bonds and binding free energy. These peptides were selected for experimental testing and were designated as peptides 1, 2 and 3 in the experimental assays. The row highlighted in grey shows the peptide that was found to be a positive hit in the agglutination assay.

| Peptide | Stable binding duration (ns) | Binding Free Energy (kcal/mol) | SEM | Protein-peptide interaction energy (kcal/mol) | Desolvation energy (kcal/mol) | Sequence | No. of stable peptide-protein hydrogen bonds |  |
| --- | --- | --- | --- | --- | --- | --- | --- | --- |
| Y36_cls3_M2_M4_R8 | 278 | -25.9 | 0.06 | -85.6 | 59.7 | ACE-AMAMGYPRASY-NH2 | 0 |  |
| **Y36_cls3_R2_M4** | **394** | **-46.2** | **0.06** | **-140.4** | **94.2** | **ACE-ARAMGYPGASY-NH2** | **2** | **Peptide 1** |
| Y36_cls5_M2_R4_W8_Y9 | 184 | -22.3 | 0.08 | -64.1 | 41.8 | ACE-AMARGYPWYSY-NH2 | 0 |  |
| **Y36_cls5_R2_F4_I8_Y9** | **500** | **-42.3** | **0.06** | **-124.5** | **82.2** | **ACE-ARAFGYPIYSY-NH2** | **1** | **Peptide 2** |
| Y45_cls1_M3_R4_M8_I10 | 172 | -16.2 | 0.06 | -66.2 | 50 | ACE-SYMRAYPMQIP-NH2 | 0 |  |
| Y45_cls1_M3_R4_M8_M10 | 91 | -35.5 | 0.2 | -87.8 | 52.3 | ACE-SYMRAYPMQMP-NH2 | 0 |  |
| Y45_cls70_Y1_M8_R9_F10_R11 | 500 | -34.4 | 0.5 | -136.6 | 102.2 | ACE-YYPGAYPMRFR-NH2 | 0 |  |
| **Y45_cls70_Y1_R8_R9_Y10_R11** | **500** | **-29.5** | **0.04** | **-169.3** | **139.8** | **ACE-YYPGAYPRRYR-NH2** | **2** | **Peptide 3** |

### Peptide testing on pre-B ALL cells

We tested these peptides in the agglutination assay. As shown in **Fig. 4A**, without treatment, LAX56 cells appear as a single-cell suspension. When GST alone was added as a negative control, (**Fig. 4B**) no agglutination was measured, whereas GST-Gal3 (**Fig. 4C**) caused cellular agglutination as expected. We used the glycomimetic (TD-139 (47, 48) and citations therein) as positive control (49) and the compound clearly inhibited Galectin-3 mediated agglutination (**Fig. 4D**). Two of the three peptides tested, P1 peptide-1 (**Fig. 4E**) and P2 peptide-2 (not shown) had no effect on agglutination. However, peptide-3 (P3) clearly was inhibitory: there was a dose-response, with a correlation between different concentrations of P3 and degree of disruption of Galectin-3-mediated lattice formation (**Fig. 4G, H; Fig. 4I**).

### Site-directed mutagenesis identifies residues important for Galectin-3 agglutination function

The model predicted strong contacts of, among others, Y36 and Y45 in the IDR with amino acids L131, L203 and H217 in the CTD (**Suppl. Fig. 2**). Therefore, the latter three amino acids were mutated to alanine to test their contribution to the Galectin-3 agglutination activity. However, GST-Gal3 L131A and GST-Gal3 L203A (**Fig. 5G and 5J, Fig. 5P** quantitation) as well GST-Gal3 H217 (not shown) still were able to agglutinate LAX56 cells, although the ability of the L131A Gal3 mutant was enhanced, and that of L203A reduced, compared to wild type Gal3 (**Fig. 5P**). Moreover, the agglutination mediated by the L203A mutant could still be inhibited by P3 peptide-3 (**Fig. 5L**), but interestingly the L131A mutant was largely insensitive to inhibition (**Fig. 5I**). We then generated a L131A/L203A double mutant. As shown in **Fig. 5M**, this mutant was functionally inactive and failed to agglutinate LAX56 leukemia cells. This identified L131 and L203 as contact points of the IDR with the shallow pocket in the CTD that are essential for agglutination.

### NMR identifies contacts of peptide-3 with the F-face of the CTD

If P3 peptide-3 inhibits GST-Gal3 mediated agglutination by interfering with the interaction of the IDR with the CTD, P3 would likely make contact with the CTD. We next used NMR to investigate this. We generated a Galectin-3 CTD construct including amino acids 117-250 and used published NMR structure data (25, 41, 50, 51) for assignments of CTD amino acid residues. As shown in **Fig. 6**, NMR showed that P3 makes extensive contacts with amino acids in the CTD. In a dose-response titration with increasing concentrations of P3, as exemplified in **Fig. 6B**, large Δδ shifts were measured with a number of amino acids such as L203, V204, A212 and L218. **Fig. 6C** provides a summary of the chemical shift perturbation (CSP) measured at the highest molar ratio of P3 to Galectin-3 CRD for the amino acid residues identified. There were seven amino acids that had a more than two-fold increased RMSD of their CSP when exposed to P3. This included residues K210, V211, A212 and V213 in the β8 sheet as well as A216, L218 and L219 in the β9 sheet. Other residues with an increased RMSD of around 2 in their CSP included V202, V204 and E205 located in the β7 sheet, and I132 in the β2 sheet. These residues are all located on the F-face of the CDR (**Suppl. Fig. 1A**). Residues such as R186, K227 or Y221 which are located in the S-face of the CRD exhibited no shift upon exposure to P3 as shown in **Fig. 6C**.

## Discussion

Proteins containing IDRs have emerged as major players in various biochemical pathways, creating unprecedented opportunities for drug targeting. However, three main challenges in designing drugs targeting IDRs using structure based approaches are (a) their lack of well-defined structure, (b) the difficulty in translating experimental structural information into three-dimensional atomic coordinates, (c) our lack of understanding how conventional drug design approaches that target specific protein structures can be applied to the ensemble of diverse conformations of an IDR (52–55). Moreover, designing therapeutics necessitates a detailed mechanistic understanding of IDR dynamics and the interaction with self or other partners.

We here used Galectin-3 as a prototypical IDR-containing protein to explore the potential of designing IDR-directed function-targeting therapeutics. Using the AMD method that efficiently samples the IDP conformational space, and existing NMR data that enables filtering the MD-derived ensemble into an experimentally relevant subset, we identified diverse NTD conformations bound to the CTD. This is unprecedented, since the NMR data alone only allowed the identification of the CTD residues that interact with the NTD, but not specific NTD structures that contribute to this interaction.

The *in silico* exploration of the Galectin-3 conformational ensemble and the subsequent mutagenesis results point to a complicated mechanistic picture of the NTD-CTD interaction. Our initial analysis of the AMD-generated ensemble pointed to two residues, Y36 and Y45 as making the most stable contacts with the CTD as well as their region of contact within the CTD. However, site-directed mutagenesis of Y36A and Y45A alone or in combination still yielded a Galectin-3 protein capable of agglutinating pre-B ALL cells (not shown). Subsequently, applying the BME approach, we obtained a more diverse structural ensemble consistent with NMR, in which, apart from Y36 and Y45, multiple IDR/NTD residues have significant interactions with the CTD. These observations are entirely consistent with Lin et al (26) who, using NTD-truncated constructs, concluded that multiple aromatic residues in the NTD interact with the CTD, and with Zhao et al (21) who mutated 14 prolines in the NTD to show that many of them are involved in the NTD-CTD interaction.

From a thermodynamic point of view, transient interactions of multiple NTD motifs with the CTD (“many-to-one” interactions) may lead to a dynamic structural ensemble with high entropy. We anticipate that this should contribute positively to the stability of the NTD-CTD complex. Stability of protein-protein interactions is determined by the free energy difference *ΔG* (which is comprised of entropic and enthalpic components) between the bound and the unbound states. In interactions involving folded proteins, the formation of the complex is associated with a large loss of configurational entropy that needs to be compensated through enthalpic gain, in order for the complex to be energetically stable (i.e. negative *ΔG*). With proteins such as Galectin-3 involving interactions with an IDR, entropic loss, relative to the unbound state in which no NTD-CTD interactions are present, can be minimized through “many-to-one” interactions. Therefore, the enthalpic gain upon binding need not be as strong as in the case of folded protein interactions. Since “many-to-one” interactions are hallmarks of many disordered proteins, the above principle applies to other IDR containing proteins with repeat motifs as well. An elegant discussion of the role of repeat motifs and entropy in IDR interaction can be found in Flock et al (56).

The AMD simulations and subsequent analysis detected a pocket in the F-face of CTD as a possible area of NTD contact and identified L131/L203 as essential for the NTD interaction. L203 was previously shown to be important for the interaction between the NTD and CTD (25) and an L203A Galectin-3 mutant has reduced capacity to form liquid-liquid phase separation droplets (21). In concordance with this, we also found that the L203A single mutant had reduced ability to agglutinate the leukemia cells although it still retained some activity.

To assess the impact of L131 and L203 on the NTD interaction, we also (**Suppl. Fig. 2**) calculated the average interaction energy of each CTD residue in the F face in the NMR filtered AMD ensemble. The top five CTD residues showing the lowest interaction energy are shown in **Suppl. Fig. 2**. We calculated the interaction energy separately in the two conformational clusters, where either Y36 or Y45 makes contact with the CTD. In both cases, L131 and L203 show up at the top, along with several other polar residues such as H217, Q201 and D215. The impact of polar residues on protein-protein interactions are likely to be small due to competition with the solvent. This leaves L131 and L203 as the key hydrophobic residues that contribute to NTD binding, with an interaction energy of −1 to −1.5 kcal/mol. According to previous computational studies on protein-protein interface (PPI) hotspots, an energy contribution > −2 kcal/mol is likely to impact binding of partner proteins significantly (57). Here, individually, the energy contributions of the two hydrophobic residues are less than −2 kcal/mol, but together, their contribution is a substantial −2 to −2.5 kcal/mol. The importance of AMD analysis was thus illustrated by the additional identification of the need for cooperativity of L131 with L203 in Galectin-3 to allow this lectin to agglutinate leukemia cells.

Although studies to inhibit Galectin-3 classically focused on blocking the binding of carbohydrate substrates to the recognition domain (14), (42), (58), galactomannins (59) and PTX008, a calixarene (50) may also target the NTD-CRD interaction. PTX008 contacts V202, K210, V211 and A216 located in the F-face of the CRD, which are also contacted by peptide-3 in our study. In contrast to peptide-3, galactomannins and PTX008 may also make some contacts with the S-face of the CTD, and inhibit Galectin-1, a lectin that has overlap in binding targets with Galectin-3 (36, 60).

Peptide-3 was identified using a completely *in s<u>ilico</u>* approach. The computational analysis reduced the large number of candidate peptides for experimental screening to a very small number. Importantly, peptide-3 includes the –PGAY- motif previously shown to be critical in the Galectin-3 NTD-CRD interaction (25). In that study, the PGAY peptide was reported to mainly contact residues G124, F192, Q201, V202, L203, V204, K210, V211, A212, V213, D215, A216, L218, L219, and Q220. According to our NMR data, the CRD residues that show significant chemical shifts in response to peptide-3 binding are I132, V202, V204, E205, K210, V211, A212, V213, A216, L218 and L219 (**Suppl. Fig. 1A and C**). According to the MD simulation, these residues are all located within 5Å of the predicted binding site of peptide-3. Thus the contacts made by the PGAY peptide and P3 have a large degree of overlap, but also some differences. However, majority of the residues that contact the PGAY peptide are located within 5Å of peptide-3, with the exception of G124 and Q220. This indicates that the gross binding sites of the two peptides are highly similar. In particular, I132, which is located in the β2 sheet of the CTD, is of interest because the adjacent mutation of L131 combined with L203 abrogated the agglutination of Galectin-3, suggesting that the β2 sheet may have a critical contribution to the IDR-CTD interaction.

It is not exactly clear how the intrinsically disordered region in Galectin-3 works together with its structured C-terminal end to achieve a biological effect such as agglutination. NMR studies have indicated that the interaction of the IDR/NTD with the CTD does not cause a strong conformational shift in critical residues of the S-face (25), ruling out a simple allosteric effect in which contact of the IDR with the F-face in the CTD would open up the carbohydrate binding site on the S-face other side and allow entry of the client glycoprotein. Zhao et al (21) showed that the IDR/NTD did not need to be physically attached to the CTD for Galectin-3 to cause liquid-liquid phase separation of a client glycoprotein, CD45, *in vitro*. Their data provide evidence for a model in which the S-face of the CTD binds to a glycoprotein and leaves the F-face to interact with the IDR of other Galectin-3 molecules as a key step in polymerization. Our BME results entirely support such a mechanism of cooperation between the NTD and the CTD by showing the involvement of multiple NTD motifs in recognizing the CTD. This can lead to stable CTD binding by minimizing entropic loss through “many-to-one” interactions, as discussed previously. Moreover, during oligomerization, when multiple CTDs from different Galectin-3 molecules could be interacting with the NTD, the repeat NTD motifs can support such interactions. In such a scenario, each NTD motif can bind to a different CTD, as opposed to multiple CTDs competing for the same NTD site for binding. Importantly, this property could allow a single NTD to interact with multiple F-face pockets in different CTDs. In this model, the switch from one NTD binding to it’s own CTD to one NTD binding to multiple CTDs could be concentration-driven, where low Galectin-3 levels may favor the former, while high protein levels the latter interactions.

## Conclusion

In the current study we have addressed the question if it is possible to interfere with the interaction of an IDR with a domain of defined structure using Galectin-3 as a test case. Our results show that this is feasible. Because IDRs are enriched in many important proteins that form RNP complexes and membrane-less subcellular compartments such as stress granules, it may be possible to use a strategy similar to the one used here to disperse complexes in which they part of and inhibit their function.

## Acknowledgements

This study was supported by NIH RO1 CA172040 to NH. We thank Yuelong Ma [Shared Resources-Synthetic Chemistry] for peptide synthesis and the Nuclear Magnetic Resonance Core for performing the NMR experiments.

## Abbreviations

AMD: accelerated molecular dynamics
CRD: carbohydrate-recognition/-binding domain
CSD: chemical shift differences
CSP: chemical shift perturbation
CTD: C-terminal domain
IDR: intrinsically disordered region
NTD: N-terminal domain
pre-B ALL: precursor B acute lymphoblastic leukemia
RNP: ribonucleotide-protein particle
RMSD: root mean square deviation

---

**Suppl. Fig. 1.**
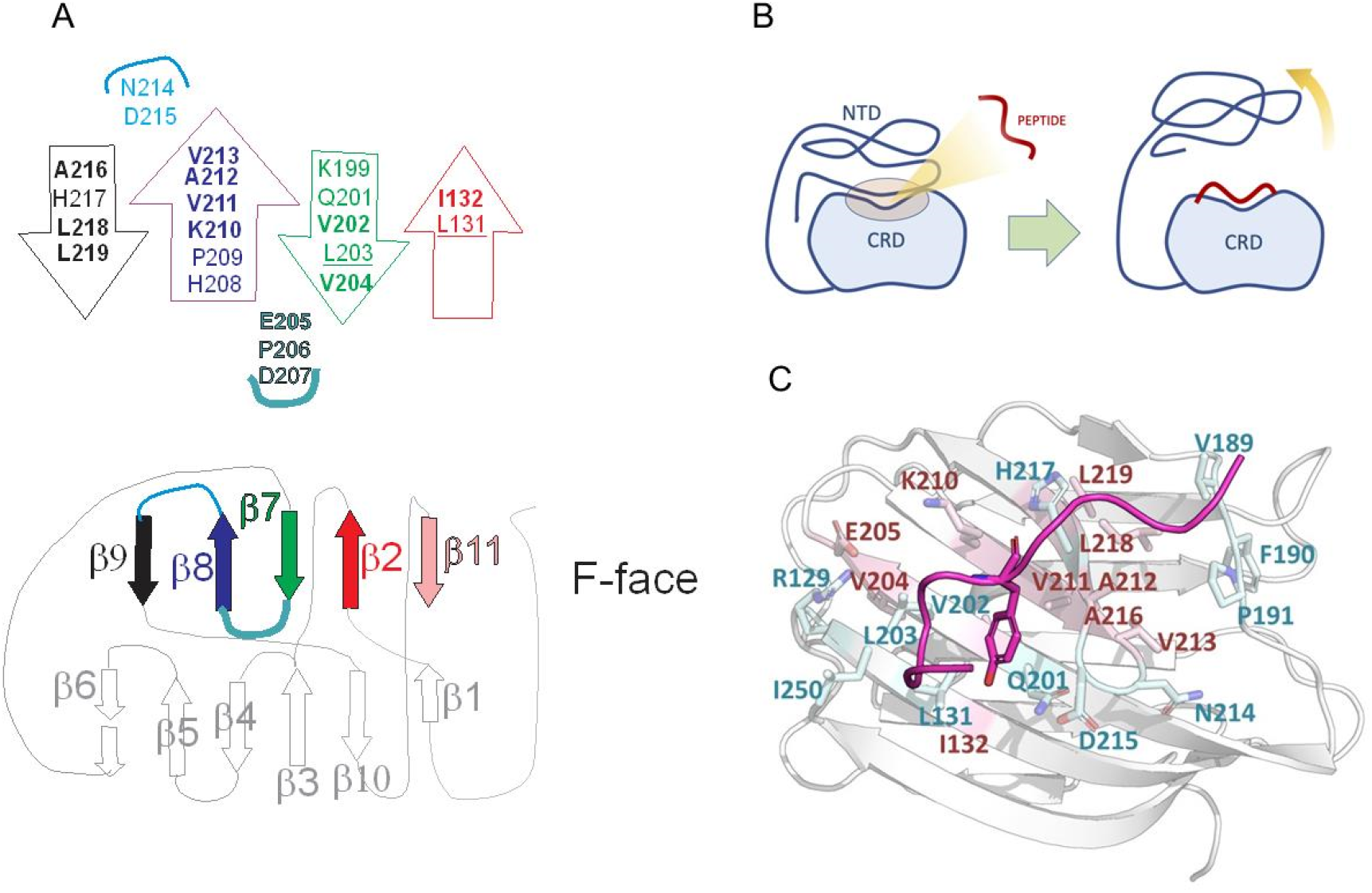
Schematic of Galectin-3. **(A)** The CTD with the β-sheets of the F-face as indicated. Amino acids present in each β sheet of the F-face are indicated. Residues making strong contact with peptide-3 are bold; L131 and L203 residues are underlined. **(B)** Peptide-3 inhibits the IDR from contacting the shallow pocket in the F-face of the CTD which can be located on the same or on a different Galectin-3 molecule. (**C)** The binding mode of peptide-3 to Galectin-3 CTD, as predicted from the MD simulation. The CTD residues within 5Å of the bound peptide are highlighted as sticks. Residues that show significant chemical shifts are highlighted in red. The peptide is shown as a magenta cartoon. The tyrosine corresponding to the central PGAY motif is displayed as a stick.

**Suppl. Fig.2.**
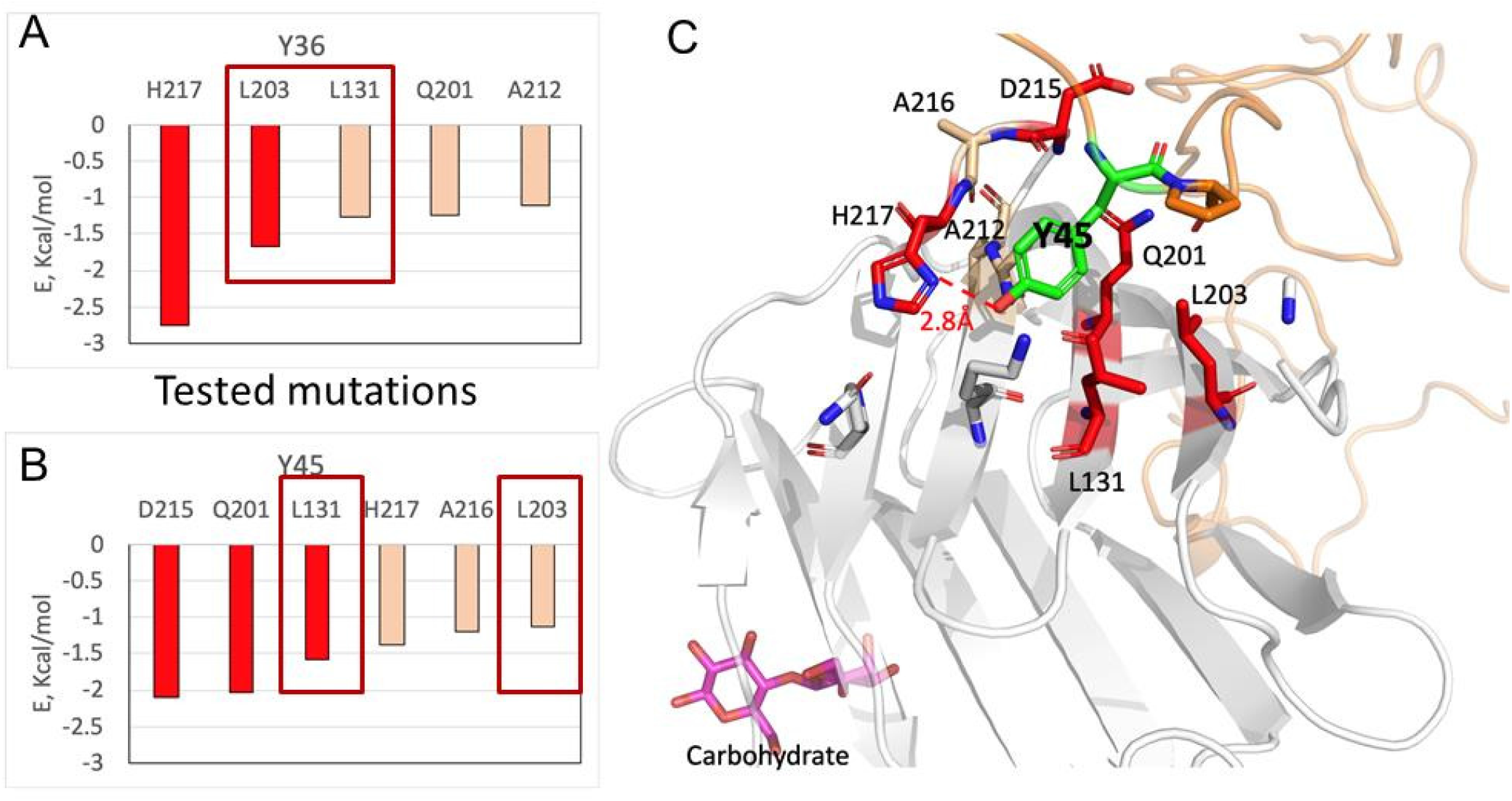
Contributions of the CTD residues in binding of NTD. **(A, B)** The interaction energies of the top CTD residues with the NTD in two major clusters derived from the AMD simulations in which either Y36 **(A)** or Y45 **(B)** of the NTD inserts into the F face. The bars representing the highest energy contribution are colored red. The residues of which mutations resulted in significant loss of agglutination are highlighted by red boxes. **(C)** Zoomed-in view of the binding pocket of the NTD in the CTD in which Y45 inserts into the F face. The CTD residues showing the highest contribution to NTD binding are highlighted in red. Y45 is colored green. The CTD and the NTD are shown as grey and orange cartoons respectively. The hydrogen bond between the -OH group of Y45 and H217 is shown as a dotted red line.

## References

1. Wright PE, Dyson HJ. Intrinsically disordered proteins in cellular signalling and regulation. Nat Rev Mol Cell Biol. 2015;16(1):18–29.

2. Afanasyeva A, Bockwoldt M, Cooney CR, Heiland I, Gossmann TI. Human long intrinsically disordered protein regions are frequent targets of positive selection. Genome Res. 2018;28(7):975–82.

3. Coppin L, Jannin A, Ait Yahya E, Thuillier C, Villenet C, Tardivel M, et al. Galectin-3 modulates epithelial cell adaptation to stress at the ER-mitochondria interface. Cell Death Dis. 2020;11(5):360.

4. Magescas J, Sengmanivong L, Viau A, Mayeux A, Dang T, Burtin M, et al. Spindle pole cohesion requires glycosylation-mediated localization of NuMA. Sci Rep. 2017;7(1):1474.

5. Jia J, Claude-Taupin A, Gu Y, Choi SW, Peters R, Bissa B, et al. Galectin-3 Coordinates a Cellular System for Lysosomal Repair and Removal. Dev Cell. 2020;52(1):69–87 e8.

6. Coppin L, Leclerc J, Vincent A, Porchet N, Pigny P. Messenger RNA Life-Cycle in Cancer Cells: Emerging Role of Conventional and Non-Conventional RNA-Binding Proteins? Int J Mol Sci. 2018;19(3).

7. Coppin L, Vincent A, Frenois F, Duchene B, Lahdaoui F, Stechly L, et al. Galectin-3 is a non-classic RNA binding protein that stabilizes the mucin MUC4 mRNA in the cytoplasm of cancer cells. Sci Rep. 2017;7:43927.

8. Joeh E, O’Leary T, Li W, Hawkins R, Hung JR, Parker CG, et al. Mapping glycan-mediated galectin-3 interactions by live cell proximity labeling. Proc Natl Acad Sci U S A. 2020;117(44):27329–38.

9. Sciacchitano S, Lavra L, Morgante A, Ulivieri A, Magi F, De Francesco GP, et al. Galectin-3: One Molecule for an Alphabet of Diseases, from A to Z. Int J Mol Sci. 2018;19(2).

10. Suthahar N, Meijers WC, Sillje HHW, Ho JE, Liu FT, de Boer RA. Galectin-3 Activation and Inhibition in Heart Failure and Cardiovascular Disease: An Update. Theranostics. 2018;8(3):593–609.

11. Farhadi SA, Liu R, Becker MW, Phelps EA, Hudalla GA. Physical tuning of galectin-3 signaling. Proc Natl Acad Sci U S A. 2021;118(19).

12. Lee JJ, Hsu YC, Li YS, Cheng SP. Galectin-3 Inhibitors Suppress Anoikis Resistance and Invasive Capacity in Thyroid Cancer Cells. Int J Endocrinol. 2021;2021:5583491.

13. Dings RPM, Miller MC, Griffin RJ, Mayo KH. Galectins as Molecular Targets for Therapeutic Intervention. Int J Mol Sci. 2018;19(3).

14. Bertuzzi S, Quintana JI, Arda A, Gimeno A, Jimenez-Barbero J. Targeting Galectins With Glycomimetics. Front Chem. 2020;8:593.

15. Blanchard H, Yu X, Collins PM, Bum-Erdene K. Galectin-3 inhibitors: a patent review (2008-present). Expert Opin Ther Pat. 2014;24(10):1053–65.

16. Stegmayr J, Zetterberg F, Carlsson MC, Huang X, Sharma G, Kahl-Knutson B, et al. Extracellular and intracellular small-molecule galectin-3 inhibitors. Sci Rep. 2019;9(1):2186.

17. Hirani N, MacKinnon AC, Nicol L, Ford P, Schambye H, Pedersen A, et al. Target inhibition of galectin-3 by inhaled TD139 in patients with idiopathic pulmonary fibrosis. Eur Respir J. 2021;57(5).

18. Bratteby K, Torkelsson E, L’Estrade ET, Peterson K, Shalgunov V, Xiong M, et al. In Vivo Veritas: (18)F-Radiolabeled Glycomimetics Allow Insights into the Pharmacological Fate of Galectin-3 Inhibitors. J Med Chem. 2020;63(2):747–55.

19. Smith BAH, Bertozzi CR. The clinical impact of glycobiology: targeting selectins, Siglecs and mammalian glycans. Nat Rev Drug Discov. 2021;20(3):217–43.

20. Dumic J, Dabelic S, Flogel M. Galectin-3: an open-ended story. Biochim Biophys Acta. 2006;1760(4):616–35.

21. Zhao Z, Xu X, Cheng H, Miller MC, He Z, Gu H, et al. Galectin-3 N-terminal tail prolines modulate cell activity and glycan-mediated oligomerization/phase separation. Proc Natl Acad Sci U S A. 2021;118(19).

22. Uchino Y, Woodward AM, Mauris J, Peterson K, Verma P, Nilsson UJ, et al. Galectin-3 is an amplifier of the interleukin-1beta-mediated inflammatory response in corneal keratinocytes. Immunology. 2018;154(3):490–9.

23. Mirandola L, Yu Y, Cannon MJ, Jenkins MR, Rahman RL, Nguyen DD, et al. Galectin-3 inhibition suppresses drug resistance, motility, invasion and angiogenic potential in ovarian cancer. Gynecol Oncol. 2014;135(3):573–9.

24. Mirandola L, Yu Y, Chui K, Jenkins MR, Cobos E, John CM, et al. Galectin-3C inhibits tumor growth and increases the anticancer activity of bortezomib in a murine model of human multiple myeloma. PLoS One. 2011;6(7):e21811.

25. Ippel H, Miller MC, Vertesy S, Zheng Y, Canada FJ, Suylen D, et al. Intra- and intermolecular interactions of human galectin-3: assessment by full-assignment-based NMR. Glycobiology. 2016;26(8):888–903.

26. Lin YH, Qiu DC, Chang WH, Yeh YQ, Jeng US, Liu FT, et al. The intrinsically disordered N-terminal domain of galectin-3 dynamically mediates multisite self-association of the protein through fuzzy interactions. J Biol Chem. 2017;292(43):17845–56.

27. Chiu YP, Sun YC, Qiu DC, Lin YH, Chen YQ, Kuo JC, et al. Liquid-liquid phase separation and extracellular multivalent interactions in the tale of galectin-3. Nat Commun. 2020;11(1):1229.

28. Flores-Ibarra A, Vértesy S, Medrano FJ, Gabius HJ, Romero A. Crystallization of a human galectin-3 variant with two ordered segments in the shortened N-terminal tail. Sci Rep. 2018;8(1):9835.

29. Eswar N, John B, Mirkovic N, Fiser A, Ilyin VA, Pieper U, et al. Tools for comparative protein structure modeling and analysis. Nucleic Acids Res. 2003;31(13):3375–80.

30. Nguyen H, Roe DR, Simmerling C. Improved Generalized Born Solvent Model Parameters for Protein Simulations. J Chem Theory Comput. 2013;9(4):2020–34.

31. Robustelli P, Piana S, Shaw DE. Developing a molecular dynamics force field for both folded and disordered protein states. Proceedings of the National Academy of Sciences. 2018;115(21):E4758–E66.

32. Hopkins CW, Le Grand S, Walker RC, Roitberg AE. Long-Time-Step Molecular Dynamics through Hydrogen Mass Repartitioning. Journal of Chemical Theory and Computation. 2015;11(4):1864–74.

33. Salomon-Ferrer R, Götz AW, Poole D, Le Grand S, Walker RC. Routine Microsecond Molecular Dynamics Simulations with AMBER on GPUs. 2. Explicit Solvent Particle Mesh Ewald. Journal of Chemical Theory and Computation. 2013;9(9):3878–88.

34. Han B, Liu YF, Ginzinger SW, Wishart DS. SHIFTX2: significantly improved protein chemical shift prediction. Journal of Biomolecular Nmr. 2011;50(1):43–57.

35. Bottaro S, Bengtsen T, Lindorff-Larsen K. Integrating Molecular Simulation and Experimental Data: A Bayesian/Maximum Entropy Reweighting Approach. In: Gáspári Z, editor. Structural Bioinformatics: Methods and Protocols. New York, NY: Springer US; 2020. p. 219–40.

36. Paz H, Joo EJ, Chou CH, Fei F, Mayo KH, Abdel-Azim H, et al. Treatment of B-cell precursor acute lymphoblastic leukemia with the Galectin-1 inhibitor PTX008. J Exp Clin Cancer Res. 2018;37(1):67.

37. George AA, Paz H, Fei F, Kirzner J, Kim YM, Heisterkamp N, et al. Phosphoflow-Based Evaluation of Mek Inhibitors as Small-Molecule Therapeutics for B-Cell Precursor Acute Lymphoblastic Leukemia. PLoS One. 2015;10(9):e0137917.

38. Fei F, Joo EJ, Tarighat SS, Schiffer I, Paz H, Fabbri M, et al. B-cell precursor acute lymphoblastic leukemia and stromal cells communicate through Galectin-3. Oncotarget. 2015;6(13):11378–94.

39. Delaglio F, Grzesiek S, Vuister GW, Zhu G, Pfeifer J, Bax A. NMRPipe: a multidimensional spectral processing system based on UNIX pipes. J Biomol Nmr. 1995;6(3):277–93.

40. Lee W, Tonelli M, Markley JL. NMRFAM-SPARKY: enhanced software for biomolecular NMR spectroscopy. Bioinformatics. 2015;31(8):1325–7.

41. Umemoto K, Leffler H. Assignment of 1H, 15N and 13C resonances of the carbohydrate recognition domain of human galectin-3. J Biomol NMR. 2001;20(1):91–2.

42. Zou J, Glinsky VV, Landon LA, Matthews L, Deutscher SL. Peptides specific to the galectin-3 carbohydrate recognition domain inhibit metastasis-associated cancer cell adhesion. Carcinogenesis. 2005;26(2):309–18.

43. Hamelberg D, Mongan J, McCammon JA. Accelerated molecular dynamics: a promising and efficient simulation method for biomolecules. J Chem Phys. 2004;120(24):11919–29.

44. Fei F, Abdel-Azim H, Lim M, Arutyunyan A, von Itzstein M, Groffen J, et al. Galectin-3 in pre-B acute lymphoblastic leukemia. Leukemia. 2013;27(12):2385–8.

45. Beard H, Cholleti A, Pearlman D, Sherman W, Loving KA. Applying physics-based scoring to calculate free energies of binding for single amino acid mutations in protein-protein complexes. PLoS One. 2013;8(12):e82849.

46. Miller BR, McGee TD, Swails JM, Homeyer N, Gohlke H, Roitberg AE. MMPBSA.py: An Efficient Program for End-State Free Energy Calculations. Journal of Chemical Theory and Computation. 2012;8(9):3314–21.

47. St-Gelais J, Denavit V, Giguere D. Efficient synthesis of a galectin inhibitor clinical candidate (TD139) using a Payne rearrangement/azidation reaction cascade. Org Biomol Chem. 2020;18(20):3903–7.

48. Chan YC, Lin HY, Tu Z, Kuo YH, Hsu SD, Lin CH. Dissecting the Structure-Activity Relationship of Galectin-Ligand Interactions. Int J Mol Sci. 2018;19(2).

49. Hsieh TJ, Lin HY, Tu Z, Lin TC, Wu SC, Tseng YY, et al. Dual thio-digalactoside-binding modes of human galectins as the structural basis for the design of potent and selective inhibitors. Sci Rep. 2016;6:29457.

50. Miller MC, Zheng Y, Suylen D, Ippel H, Canada FJ, Berbis MA, et al. Targeting the CRD F-face of Human Galectin-3 and Allosterically Modulating Glycan Binding by Angiostatic PTX008 and a Structurally Optimized Derivative. ChemMedChem. 2021;16(4):713–23.

51. Zhang Z, Miller MC, Xu X, Song C, Zhang F, Zheng Y, et al. NMR-based insight into galectin-3 binding to endothelial cell adhesion molecule CD146: Evidence for noncanonical interactions with the lectin’s CRD beta-sandwich F-face. Glycobiology. 2019;29(8):608–18.

52. Bhattacharya S, Lin X. Recent Advances in Computational Protocols Addressing Intrinsically Disordered Proteins. Biomolecules. 2019;9(4).

53. Joshi P, Vendruscolo M. Druggability of Intrinsically Disordered Proteins. Adv Exp Med Biol. 2015;870:383–400.

54. Uversky VN. Intrinsically Disordered Proteins. Structural Biology in Drug Discovery2020. p. 587–612.

55. Cheng Y, LeGall T, Oldfield CJ, Mueller JP, Van YY, Romero P, et al. Rational drug design via intrinsically disordered protein. Trends Biotechnol. 2006;24(10):435–42.

56. Flock T, Weatheritt RJ, Latysheva NS, Babu MM. Controlling entropy to tune the functions of intrinsically disordered regions. Current Opinion in Structural Biology. 2014;26:62–72.

57. Zerbe BS, Hall DR, Vajda S, Whitty A, Kozakov D. Relationship between hot spot residues and ligand binding hot spots in protein-protein interfaces. Journal of chemical information and modeling. 2012;52(8):2236–44.

58. Stasenko M, Smith E, Yeku O, Park KJ, Laster I, Lee K, et al. Targeting galectin-3 with a high-affinity antibody for inhibition of high-grade serous ovarian cancer and other MUC16/CA-125-expressing malignancies. Sci Rep. 2021;11(1):3718.

59. Miller MC, Ippel H, Suylen D, Klyosov AA, Traber PG, Hackeng T, et al. Binding of polysaccharides to human galectin-3 at a noncanonical site in its carbohydrate recognition domain. Glycobiology. 2016;26(1):88–99.

60. Miller MC, Klyosov AA, Mayo KH. Structural features for alpha-galactomannan binding to galectin-1. Glycobiology. 2012;22(4):543–51.

